# Atomistic Mechanism of Cation Permeation and Pore Voltage Sensitivity in Cyclic Nucleotide-Gated CNGA1 Ion Channel

**DOI:** 10.1101/2024.12.19.629380

**Authors:** Haoran Liu, Johann Biedermann, Han Sun

**Affiliations:** Research Unit of Structural Chemistry & Computational Biophysics, Leibniz-Forschungsinstitut für Molekulare Pharmakologie, Berlin 13125, Germany; Department of Chemistry, Technische Universität Berlin, Berlin 10623, Germany

## Abstract

Mammalian cyclic nucleotide-gated (CNG) ion channels play a fundamental role in signal transduction within the visual and olfactory sensory cells, converting external stimuli into electrical signals. Here, using large-scale atomistic molecular dynamics (MD) simulations under transmembrane voltages, we uncover the atomistic mechanism of monovalent cation permeation in the homotetrameric CNGA1 channel, involving hydrated cations in the selectivity filter. We propose that hydration fluctuations in the central gate region, driven by pore flexibility, are the underlying mechanism for excess single-channel conductance noise and characteristic single-channel flickering. Furthermore, we suggest an atomistic mechanism for intrinsic voltage sensitivity in the channel pore, mediated by allosteric coupling between the selectivity filter and the central cavity gate. Our study provides atomistic insights into non-selective cation permeation and voltage sensitivity of the CNGA1 channel pore that could not be resolved by static structural analysis alone.

## Introduction

Cyclic nucleotide-gated (CNG) ion channels are present in both prokaryotic and eukaryotic genomes. In eukaryotes, they are found in photoreceptors, olfactory sensory neurons, and the central nervous system^1, 2, 3, 4^. In response to changes in intracellular concentrations of cAMP or cGMP, which are linked to various biochemical signaling pathways, CNG channels open or close, regulating the flow of cations across the cell membrane^5, 6^. Mutations in CNG channels are associated with a variety of channelopathies. For example, mutations in the A1 subunit of human cone CNG channels cause retinitis pigmentosa, characterized by progressive degeneration of rod and cone receptors, leading to partial or full blindness^7, 8^.

CNG channels are non-selective cation channels that do not discriminate among monovalent cations and even allow the permeation of divalent cations such as Ca^2+^. However, the opening time of the channel is largely influenced by the presence of different cation types, with longer opening time for K^+^ than Na^+ 9,10^. At the single-channel level, Na^+^ revealed approximately 1.8 times larger unitary conductance compared to K^+^. Furthermore, both Na^+^ and K^+^ showed a wider conductance distribution in the open state compared to the closed one, which was hypothesized to be caused by pore fluctuation^10^. Although CNG channels possess a voltage-sensing domain (VSD), they display little voltage sensitivity in their macroscopic currents, which is in strong contrast to their structural homologues, hyperpolarization-activated cyclic nucleotide-gated (HCN) channels^11, 12, 13, 14^. However, voltage sensitivity and gating have been observed for CNG channels in the presence of Rb^+^, Cs^+^, and organic cations^15^ as well as upon mutation in the pore^16^, under low pH conditions^17^, or at low cGMP concentration^18^. Although mutagenesis studies have revealed that voltage dependency involves residues in the pore^15, 16, 19^, the underlying atomistic mechanism remains unknown.

Structurally, CNG channels exist as both homo-and heterotetramers, with native human CNG channels forming heteromers consisting of homologous alpha (A) and beta (B) subunits^20, 21, 22^. In rod photoreceptor, the native CNG channel consists of three A1 subunits and one B1 subunit, while the cone CNG channel is formed by A3 and B3 subunits in a stoichiometry of either 3:1 or 2:2. In olfactory sensory neurons, the native CNG comprises two A2 subunits, one A4 subunit, and one B1b subunit^23, 24, 25, 26, 27, 28^. Each A and B subunit includes a pore domain, a voltage-sensing domain (VSD), a C-terminal linker (C-linker), and a cyclic nucleotide-binding domain (CNBD) (Fig. 1a). Four residues in the pore domain, T362, I363, G364, and E365, form the selectivity filter (SF), which allows only cations but not anions to pass through the channel (Fig. 1c). The first atomistic structures of the apo closed and cGMP-bound open states of CNG channels were resolved for TAX-4, a homomeric CNG channel present in *C.elegans*^29,30^. Later, structures of human homomeric CNGA1, heteromeric CNGA1/B1 rod channel, and the cone CNGA3/B3 channel in the ligand-bound open and apo closed states were resolved^27, 31, 32, 33^. These structures revealed an unprecedented but conserved gating mechanism among different CNG channels governed by a single central cavity gate involving two hydrophobic residues, for example F389 and V393 in case of CNGA1 (Fig. 1c). The SF conformations do not differ between the open and closed states of these CNG channels, resolving a long-standing debate about whether the SF functions as a gate, similar to some potassium channels^34,35^.

**Fig. 1.**
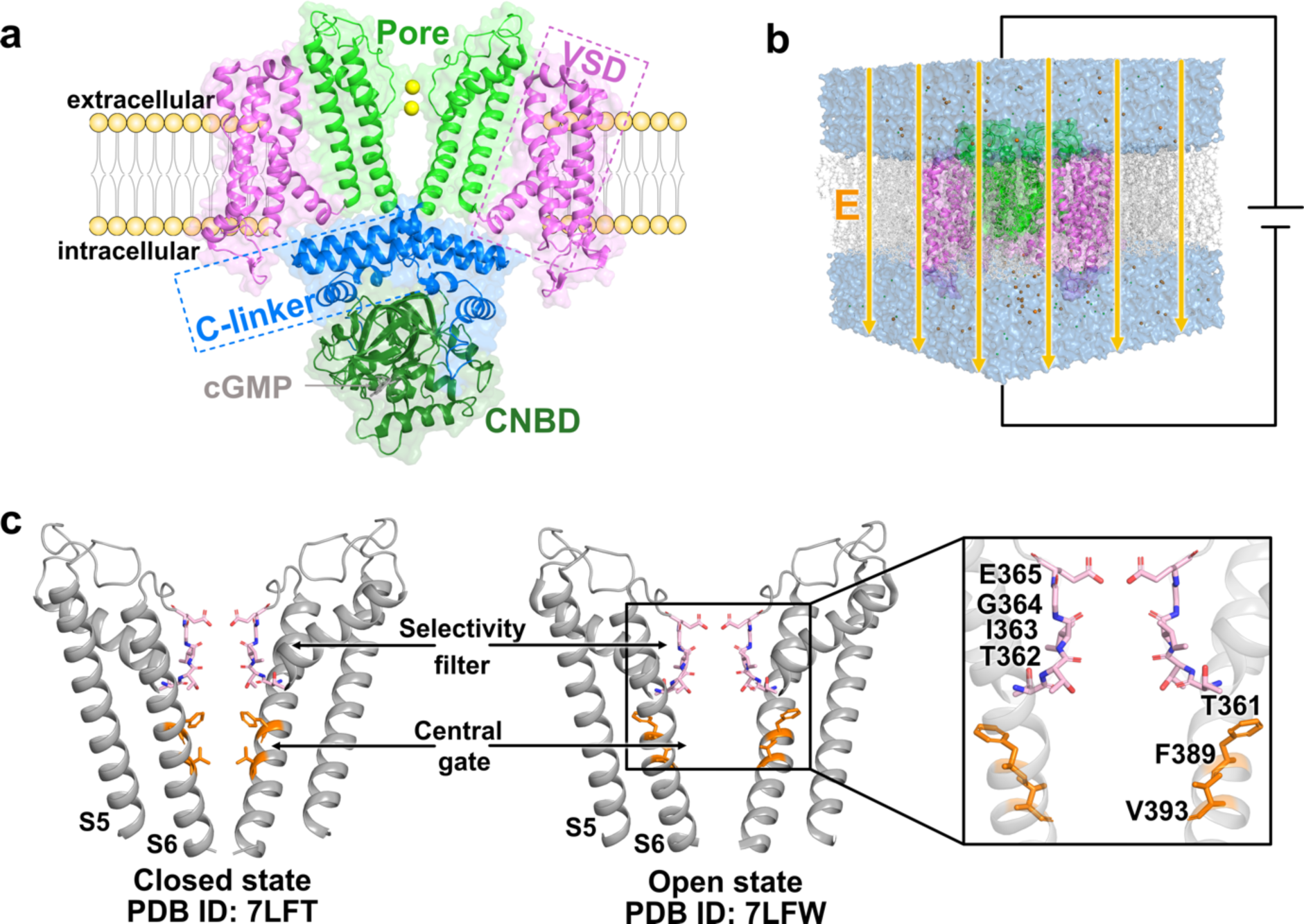
Overview of CNGA1 channel structure and MD simulation setup. **a** Structure of the cGMP-bound CNGA1 channel (PDB ID: 7LFW)^32^ within the membrane. The channel is a tetramer composed of four subunits, including the pore domain (green), voltage-sensor domain (VSD, pink), C-linker (blue), and cyclic nucleotide-binding domain (CNBD, deep green), with cGMP depicted as gray sticks. For clarity, only two diagonally opposed subunits are shown. **b** The transmembrane potential during the MD simulation is established by applying an external electric field across the simulation box. **c** Structural details of the pore domain in both the open (PDB ID: 7LFW) and closed (PDB ID: 7LFT) conformations^32^. Key residues in the selectivity filter (pink) and central gate residues (orange) are highlighted and labeled.

Based on the first open and closed structures of human CNGA1 channels, we investigated here the atomistic mechanisms of ion permeation and voltage sensitivity in CNGA1 by performing the atomistic molecular dynamics (MD) simulations under transmembrane voltages^36^ (Fig. 1b). In recent years, MD simulations have become an indispensable theoretical approach, bridging high-resolution structural analysis and electrophysiological characterization of ion channels. These computational studies together with the experimental validation enabled a detailed understanding of the atomistic mechanisms of ion permeation, gating, pharmacological modulation, and post-transcriptional regulation of various ion channels^37, 38, 39^. In this study, we demonstrated that non-selective permeation of Na^+^ and K^+^ in the CNGA1 channel involves three main ion-binding sites within and around the SF region, with partial dehydration of the cations’ first water shell at their major ion-binding sites. Furthermore, we showed that hydration fluctuation at the central cavity gate highly influences the ion conductance level, providing an atomistic understanding of the excess open state single-channel conductance noise and rapid gating characteristic. Additionally, we uncovered an unprecedented voltage sensitivity in the CNG channel pore, which might link to the high voltage sensitivity observed in CNG channels under certain conditions. This finding offers new insights for studying and designing voltage sensitivity in other ion channels and membrane proteins.

## Results

### Main ion binding profiles during ion permeation

We performed a large number of atomistic MD simulations starting from the open (PDB ID: 7LFW) and closed states (PDB ID: 7LFT) of the CNGA1 channel^32^ with K^+^ and Na^+^, and under a range of positive and negative voltages (Supplementary Table S1). These simulations included either only the pore domain or both the pore domain and the VSD. The CNGA1 channel structures were embedded within a palmitoyloleoyl phosphatidylcholine (POPC) lipid bilayer and subjected to an external electric potential to establish the transmembrane voltages^36^. The simulations were mainly performed using the CHARMM36m force field^40^, while several simulations were conducted with the AMBER99SB^41^ and AMBER19SB^42^ force fields for validation purposes (Supplementary Table S1).

From the simulations, we observed a large number of K^+^ and Na^+^ permeation events in the open state of the CNGA1 channel during the simulated timescale, while no permeation occurred in the closed state simulations on the same duration (Fig. 2a; exemplary ion permeation events for K^+^ and Na^+^ in CNGA1 are shown in Supplementary Movie S1 and S2, respectively). Along the ion conduction pathway in the open state, cation occupation was primarily observed within the SF region, with very low ion occupancy in the central cavity region. When analyzing the ion binding profiles during different simulation setups, we noted no substantial voltage-dependent variation in ion occupancy, and Na^+^ and K^+^ exhibited similar ion occupation within and around the SF region (Fig. 2a, b).

**Fig. 2.**
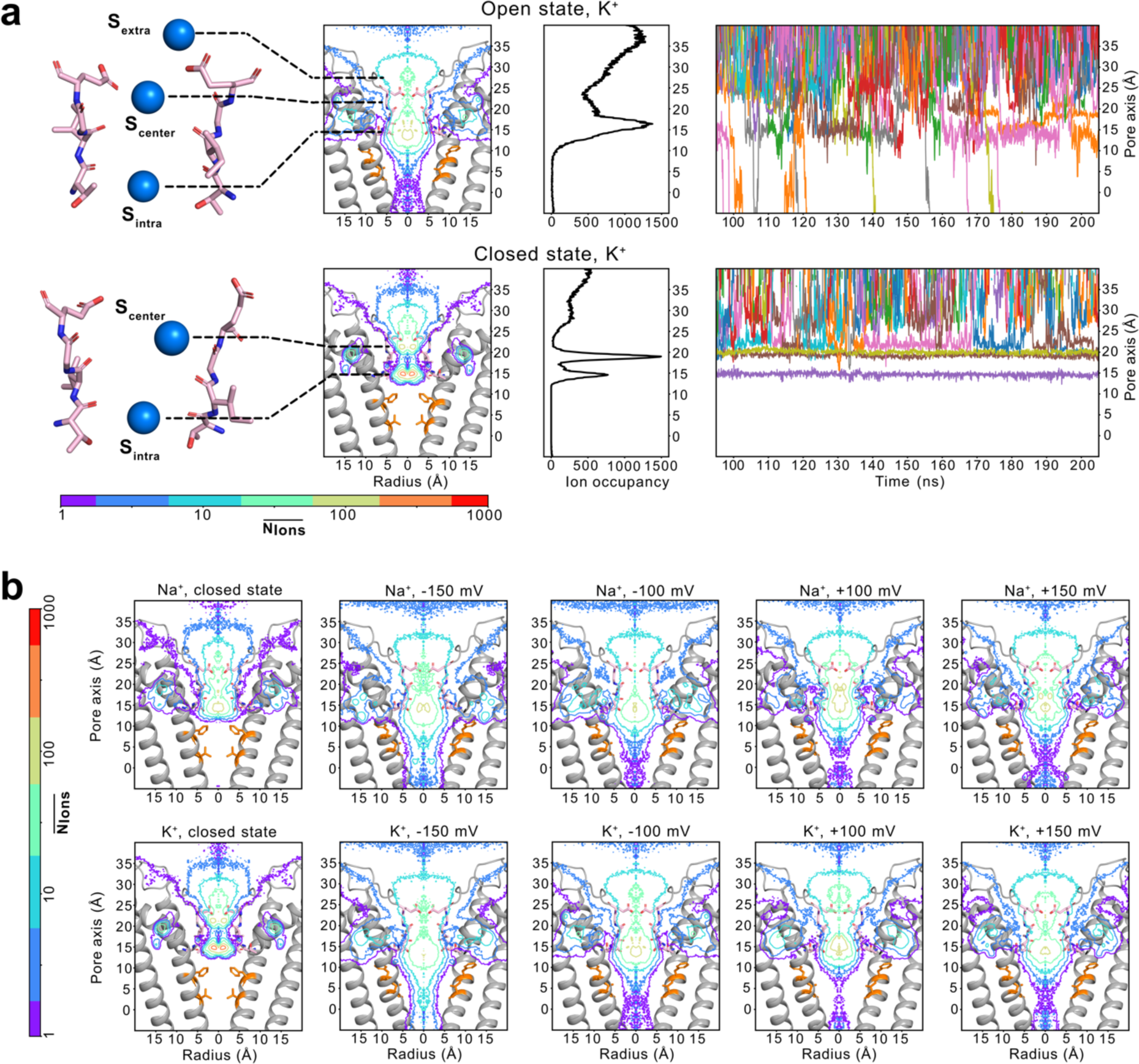
Ion binding profiles of K^+^ and Na^+^ within the ion conduction pathway derived from atomistic MD simulations. **a** (1^st^ row) major cation-binding sites within the SF; (2^nd^ row) 2-dimensional K^+^ ion occupancy maps along the pore; (3^rd^ row) 1-dimensional K^+^ ion occupancy profiles along the pore; and (4^th^ row). Representative traces showing K^+^ ions passing through the pore of the CNGA1 channel in the (top) open state and (bottom) closed state CNGA1 channel. **b** Comparison of 2-dimensional ion occupancy maps for the CNGA1 channel pore in closed (1^st^ row) and open states (2-4 row), under varying voltage conditions, with Na^+^ and K^+^.

We conducted a more detailed examination of the ion-binding profiles in the SF of the CNGA1 channel, based on MD simulations, and identified three major ion-binding regions: (i) at the lower end of the SF, coordinated by the backbone carbonyls and/or side chains of T362 residues, denoted as S_intra_, which closely corresponds to *site 2* in the previous cryo-EM analysis of CNGA1 channel^32^; (ii) in the central part of the SF, surrounded by the backbone carbonyls of I363 and partially by the side chains of E365, denoted as S_central_, which corresponds to *site 1* in the cryo-EM analysis^32^; and (iii) at the extracellular entrance of the filter, coordinated by the side chains of the negatively charged E365, denoted as S_extra_ (Fig. 2a). The electron density for this binding site was very weak in the cryo-EM map and was not further analyzed^32^, which is rather unexpected given the acidic nature of E365. From the MD simulations, we found that the binding at the extracellular entrance is pronounced for both K^+^ and Na^+^ under both negative and positive transmembrane potentials (Fig. 2b and Supplementary Fig. S1). However, this binding site is more dynamic compared to the binding sites within the SF due to the high mobility of the E365 side chain (Supplementary Fig. S2). Therefore, we hypothesize that the absence of this binding site in the cryo-EM analysis is likely due to its dynamic nature. In general, K^+^ and Na^+^ binding with the SF region of the CNGA1 channel are more diffusive and mobile compared to that in potassium and HCN channels^32, 43, 44, 45, 46, 47, 48^, most probably due to the wide dimension of the SF.

### Hydration state of ions during permeation

Next, we characterized the hydration state of K^+^ and Na^+^ ions during their permeation. Due to the considerably wider dimension of the SF of CNG channels compared to that of K^+^ channels, it was hypothesized that the cations pass through the CNG filter in a partially hydrated form^32, 49^, which is in strong contrast to K^+^ channels and also to NaK, a non-selective cation channel considered to be a bacterial homologue of CNG and HCN channels^50, 51, 52^. The analysis of our MD simulations aligned well with this hypothesis and showed that both K^+^ and Na^+^ become partially dehydrated in the filter of the CNGA1 channel, exhibiting highly comparable hydration profiles for K^+^ and Na^+^ (Fig. 3a, b). At their major binding sites in the SF, cations lose a maximum of three water molecules, which are replaced by coordination with two to three protein oxygens. At the central cavity gate, cations always retain their full hydration state during permeation.

**Fig. 3.**
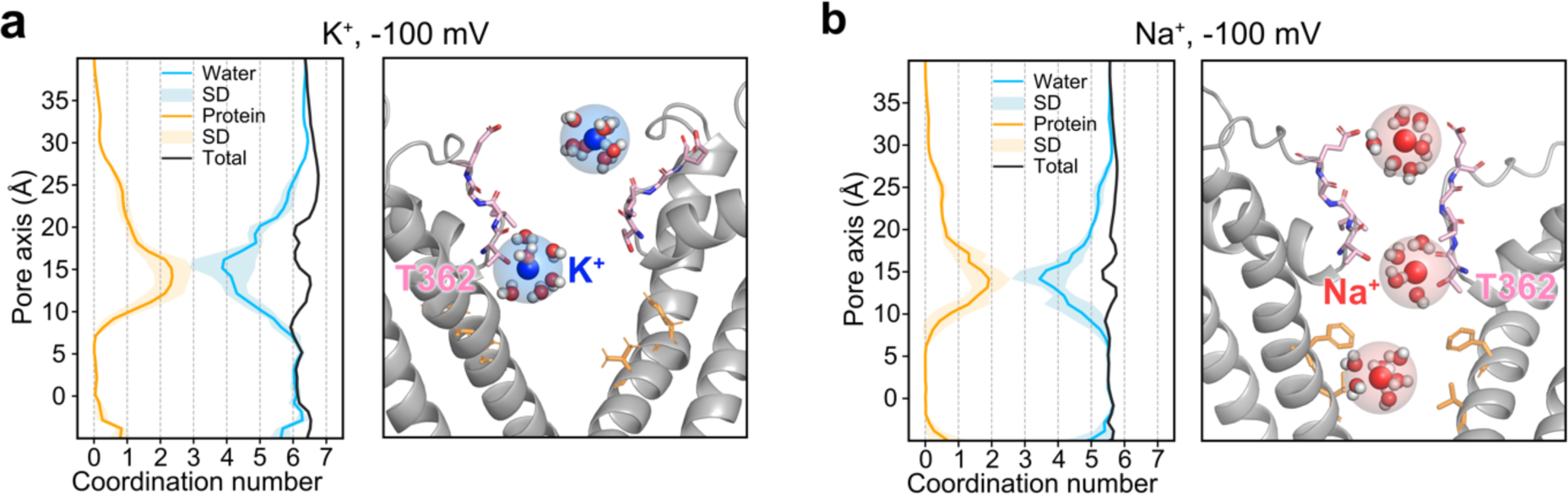
Hydration profiles of Na^+^ and K^+^ during their permeation in CNGA1 channel. **a, b** (left) Ion hydration profiles of K^+^ and Na^+^ respectively. The orange line represents the protein oxygens within the first hydration shell of the cation and the light orange shading indicates the standard deviation. The blue line represents the water oxygens within the first hydration shell and the light blue shading indicates the standard deviation. The black line represents the total number of oxygens within the first hydration shell of the ion. The snapshot (right) shows the representative snapshots selected from K^+^ (blue) and Na^+^ (red) simulations, respectively. The first hydration shell is represented as a transparent sphere.

### Comparison of simulated and experimental conductance

In this study, we simulated ion permeation in the CNGA1 channel using two different constructs: (i) the complete transmembrane domain, which includes the pore domain and the VSD (Fig. 4a); and (ii) the pore domain only (Fig. 4b, c), a common approach in previous MD studies of other ion channels to reduce the computational demand^53, 54, 55^. When comparing the simulated conductance from simulations including both the pore domain and the VSD, Na^+^ exhibited a significantly larger conductance than K^+^, along with substantially larger fluctuations among different simulation runs for Na^+^ (Fig. 4a). Generally, the simulated K^+^ conductance at physiologically relevant transmembrane voltages (-100 mV and 100 mV) falls within the range of experimental single-channel conductance of 36.0 ± 1.7 pS for Na^+^ and 20.0 ± 1.3 pS for K^+^, measured at +100 mV and at saturating cGMP concentration^10^. Simulated K^+^ conductance at -200 mV revealed irregular high conductance with large fluctuations across different simulation runs, which might be caused by large pore fluctuation due to unphysiological transmembrane voltages. Notably, the simulated Na^+^ conductance is significantly higher than the experimental value but qualitatively matches the experimental trend, which also shows larger Na^+^ conductance compared to K^+^.

**Fig. 4.**
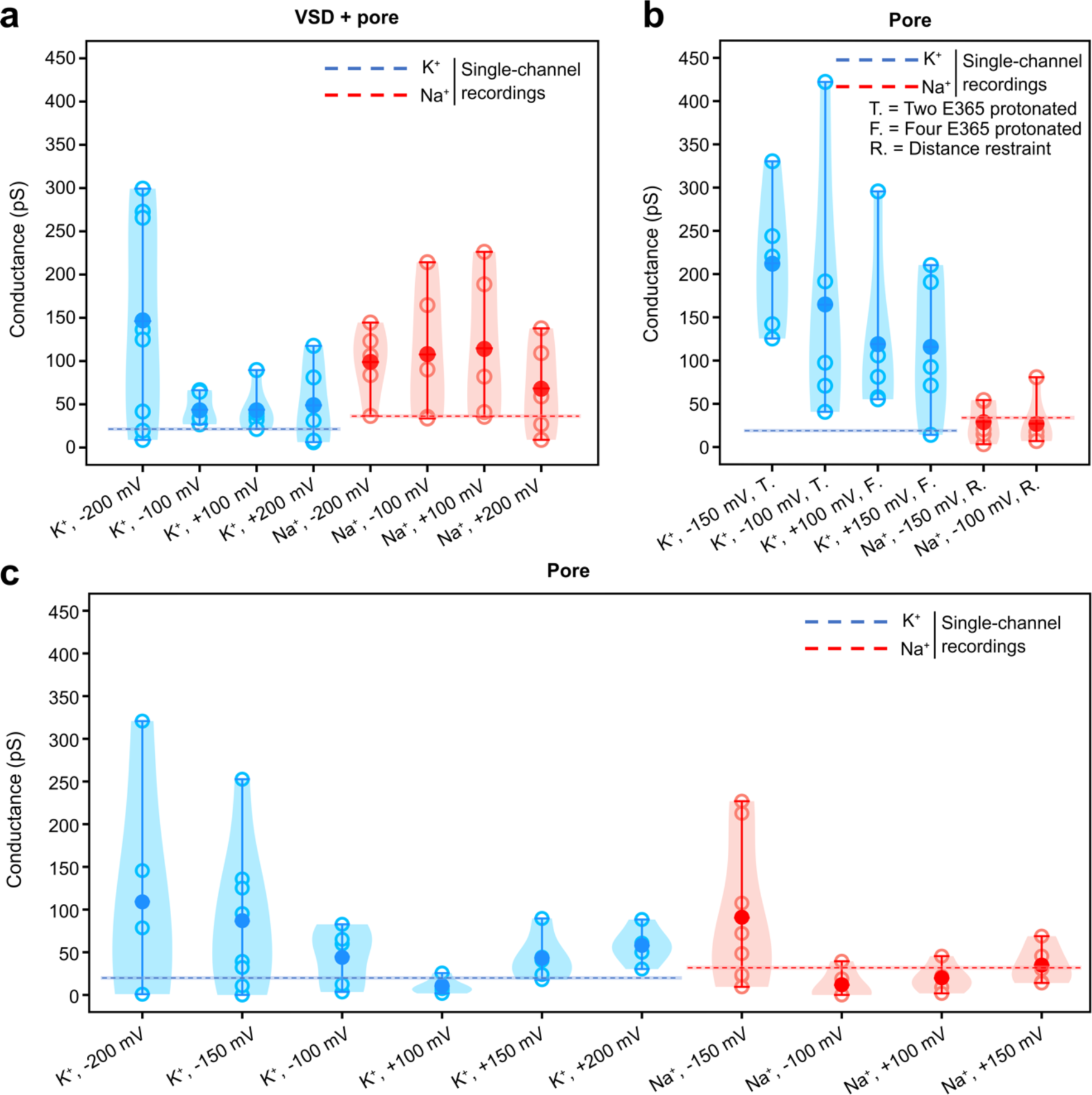
Comparison of conductance across different simulation setups. **a** Conductances derived from simulations of the pore domain and the voltage-sensor domain (VSD) of the CNGA1 channel with K^+^ (blue) and Na^+^ (red) under various transmembrane voltages. **b** Conductances derived from simulations of the pore domain of the CNGA1 channel with various configurations. “T.” represents simulations where two opposing E365 residues are protonated. “F.” represents the simulations where all four E365 residues are protonated. “R.” represents simulations with distance restraints on gate residues F389, based on distances of the open state in the cryo-EM structures^32^. **c** Conductances derived from simulations of only pore domain of the CNGA1 channel with K^+^ (blue) and Na^+^ (red) under different transmembrane voltages. Each open circle represents the conductance from a single simulation run, while the solid dots represent the average conductance from five parallel simulation runs. Conductances from experimental single-channel recordings for K^+^ (blue, 20.0 ± 1.3 pS, +100 mV) and Na^+^ (red, 36.0 ± 1.7 pS, +100 mV) from a previous study^10^ are shown as dashed lines, with error margins represented by shaded areas.

We next investigated why Na^+^ exhibits a larger single-channel conductance than K^+^ in the MD simulations of the pore domain and the VSD. It was previously proposed that Na^+^ has a shorter dwell time in the pore compared to K^+10^. To test this, we calculated the dwell times of K^+^ and Na^+^ traversing the pore from ion permeation simulations. Contrary to the initial proposal, our simulations indicated that Na^+^ had a longer dwell time in the pore than K^+^ in all simulations with different transmembrane voltages (Fig. 5a). To further understand this discrepancy, we calculated the average number of ions present in the SF and the entire pore region for both Na^+^ and K^+^. The simulations consistently showed a significantly higher average number of Na^+^ in both the SF and the pore region compared to K^+^ (Fig. 5b). Based on these MD results, we propose that the increased density of Na^+^ ions in the pore compared to K^+^, rather than the previously presumed shorter dwell time, is the primary reason for the higher single-channel conductance of Na^+^.

**Fig. 5.**
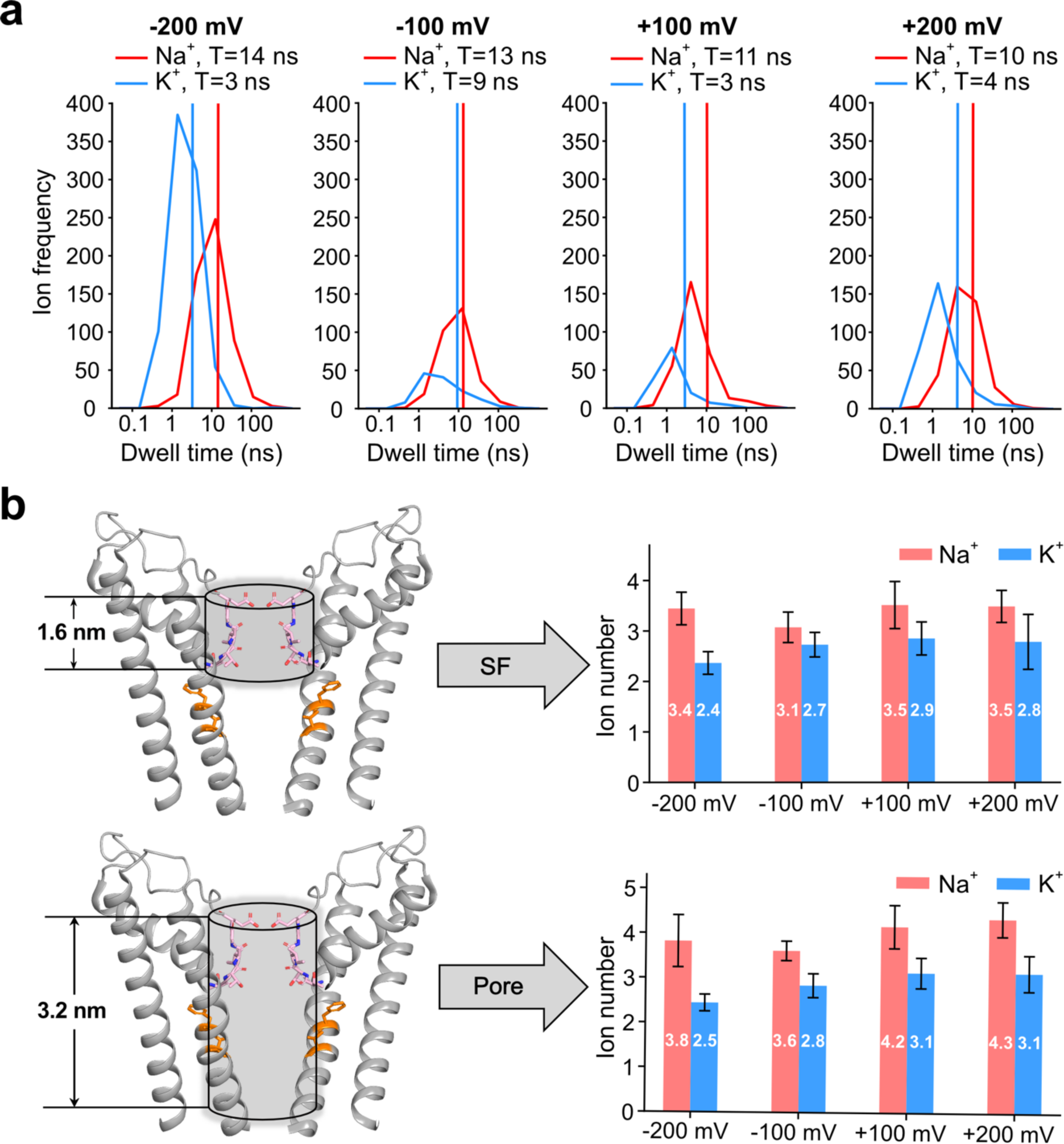
Ion dwell time and density in the pore. **a** Distribution of dwell times for Na^+^ (red) and K^+^ (blue) ions traversing the pore at various transmembrane voltages, with vertical lines indicating the means of the respective dwell times. **b** Average number of Na^+^ (red) and K^+^ (blue) in the SF and pore at different voltages. Simulations were performed with the pore domain and VSD.

### Ion conductance influenced by hydration fluctuation at the central cavity gate

As discussed earlier, we observed significant fluctuations in ion conductance across different simulation runs (Fig. 4a). Upon examining individual trajectories, we noted that in some simulations, ion permeation ceased after a certain period, while in others the channel remained conductive throughout the entire 900 ns simulation (Supplementary Fig. S3). Notably, ion permeation often stopped when the gate region became partially dehydrated, even if the channel had not fully transitioned to the closed state (Supplementary Fig. S3). To quantify this behavior, we plotted the number of ion permeation events per 20 ns simulation interval against the average number of water molecules in the gate region (F389 and V393) (Fig. 6b) within the same interval for both K^+^ and Na^+^ simulations under -100 mV (Fig. 6c) and other voltages (Supplementary Fig. S4, S5). This analysis revealed a clear relationship: the number of ion permeation events is directly correlated with the number of water molecules in the gate region.

**Fig. 6.**
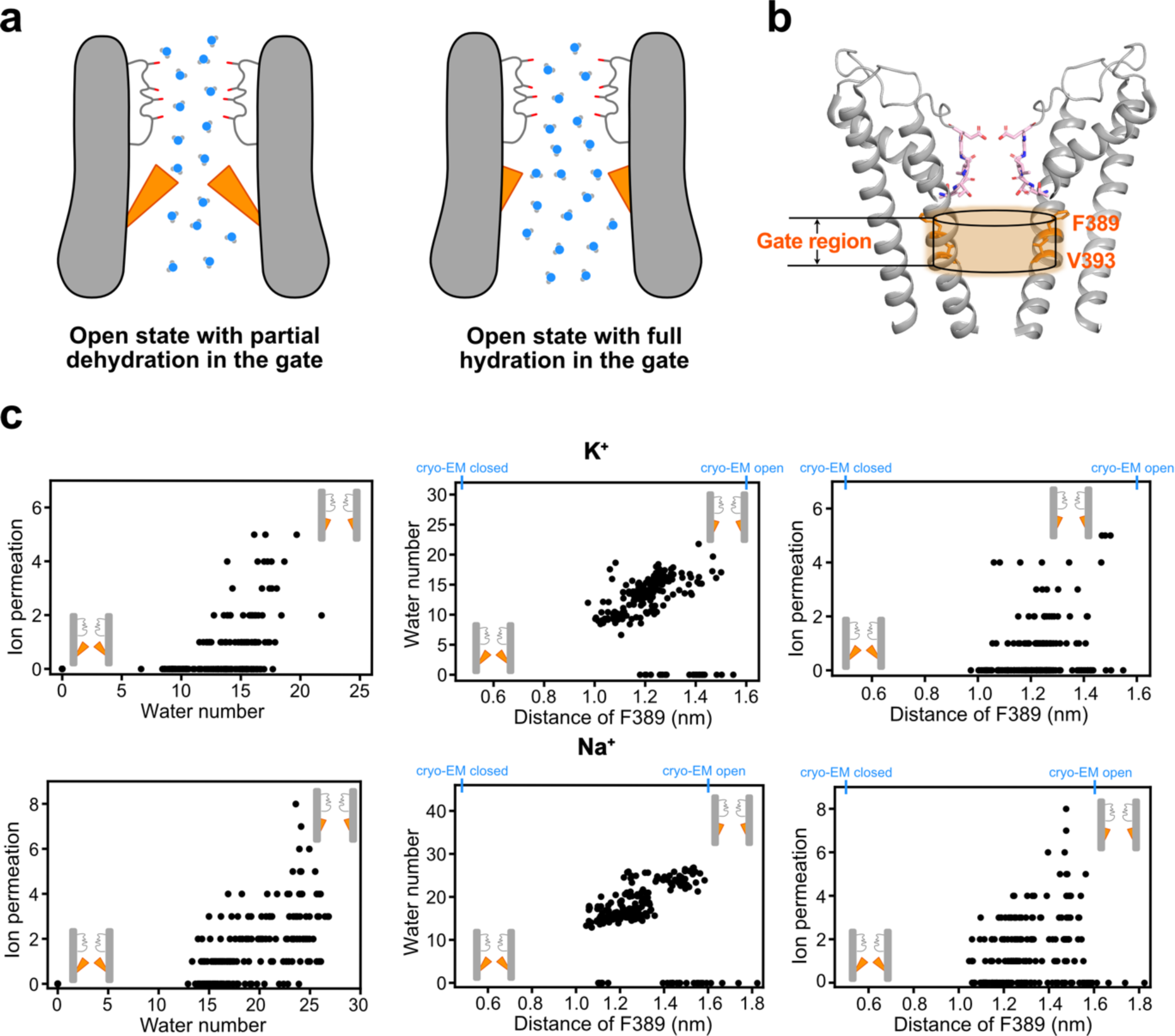
Dehydration and hydrophobic restriction at the central gate. **a** A model illustrating the CNGA1 channel in the open state, showing (left) partial dehydration and (right) full hydration in the gate region. **b** The gate region (F389 and V393) is shown as an orange cylinder. **c** The relationship between the average number of water molecules in the gate region per 20 ns interval, the average distance between two opposing gate residues (F389), and the number of ion permeation events during the same time interval. The distances between two opposing gate residue (F389) from the closed (0.5 nm) and open (1.6 nm) cryo-EM structures of CNGA1 channel^32^ are shown at the top of the figures. Simulations were performed for (top) K^+^ and (bottom) Na^+^ under -100 mV.

Furthermore, we plotted the average minimum distance between the two opposing gate residues, F389, within each 20 ns simulation interval against the number of water molecules in the gate region during the same intervals (Fig. 6c). For both K^+^ and Na^+^ under -100 mV, the analysis consistently showed a positive correlation between these two quantities. Together, our data suggest that the two large hydrophobic residues, F389 and V393, form a hydrophobic restriction site for ion permeation. Their fluctuations from the open state toward the closed state lead to changes in the hydration level of the gate region, thereby regulating ion conductance. Importantly, the channel does not need to reach a fully closed state to become non-conductive. This finding offers a plausible explanation for the excess open-channel noise and characteristic single-channel flickering of CNGA1 channels observed experimentally^10^.

### Voltage sensitivity of the pore

As mentioned above, we also simulated K^+^ and Na^+^ permeation through the CNGA1 channel by including only the pore domain. In contrast to simulations that included both the pore and the VSD, we observed notable voltage sensitivity in the simulated conductance, which increased with rising voltages (Fig. 4c). This trend was consistent for both K^+^ and Na^+^ under both positive and negative voltages. Since we found that hydration changes caused by the fluctuations in the gate residues strongly influenced the conductance, we performed two additional simulations at -100 mV and -150 mV, applying position restraints at F389 to keep the gate residues in their initial cryo-EM open state structure^32^ throughout the simulations. These new simulations, with restrained gate residues, revealed diminished voltage sensitivity (Fig. 4b).

To further understand the impact of voltage on the gate residues and the conductance, we compared the distributions of the minimum distance between the two opposing F389 residues from simulations at -100 mV, -150 mV, and -200 mV (Fig. 7a). Additionally, we plotted the average minimum distance between the opposing F389 gate residues within each 20 ns simulation interval against the number of ion permeation events in the same time interval (Fig. 7b). This analysis indicated a larger gate distance sampled at higher transmembrane voltages, which corresponded to an increased number of ion permeation events.

**Fig. 7.**
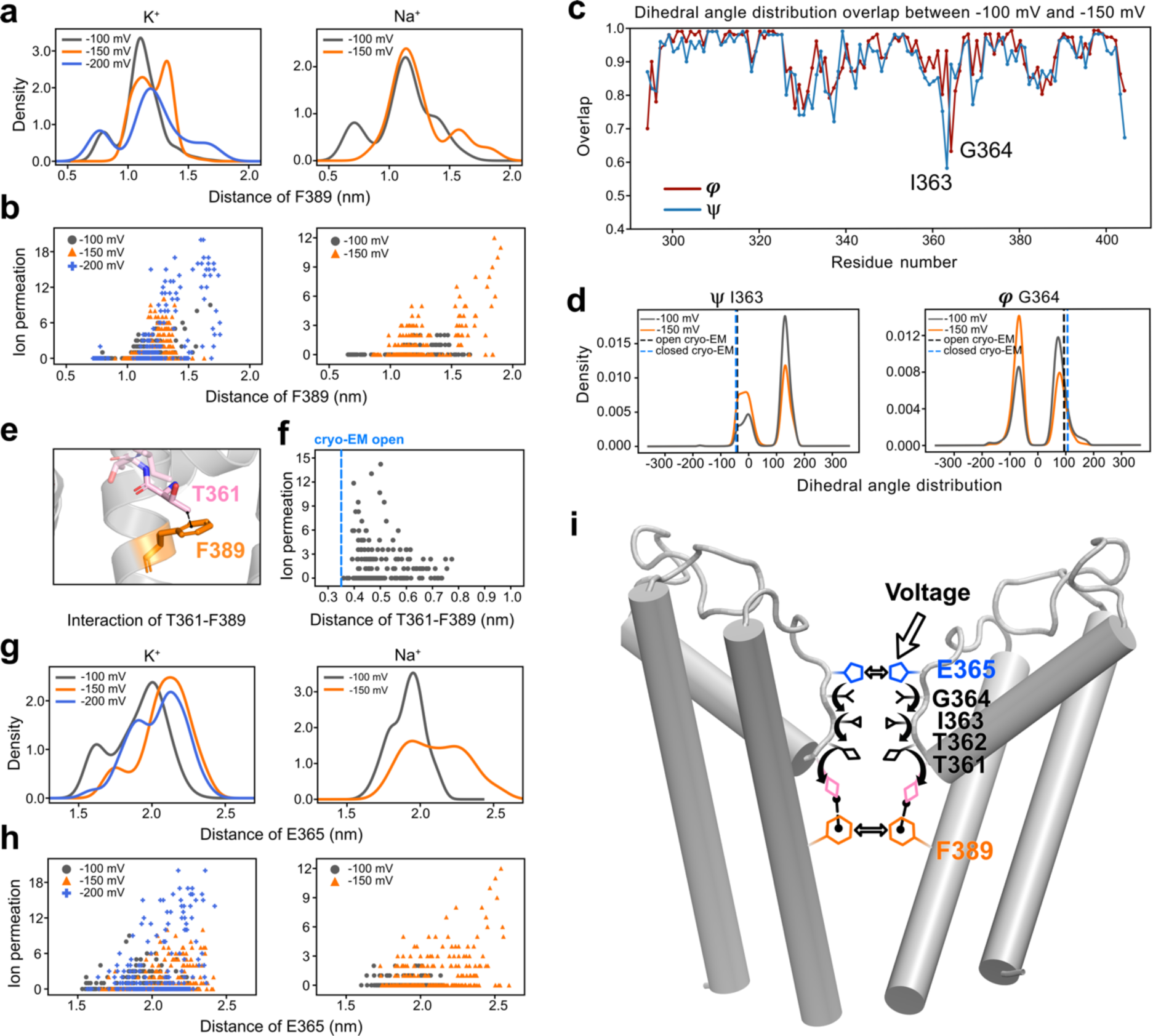
Voltage-dependent conformational changes within and surrounding the SF in simulations of the CNGA1 channel pore. **a** Distance distribution of F389 at three different voltages for K^+^ and Na^+^. **b** The relationship between the number of ion permeation per 20 ns interval and the corresponding average minimum distance between opposing F389 are shown for three different voltages. **c** Backbone dihedral angle distribution overlap between simulations at -100 mV and -150 mV for each residue in CNGA1 channel. **d** ψ angle distribution of I363 and *φ* angle distribution of G364 at -100 mV (gray) and -150 mV (orange). The dashed line represents their angles in cryo-EM structures. **e** Interaction between T361 and F389, mediated by a methyl-*π* interaction. **f** The relationship between the number of ion permeation per 20 ns interval and the average minimum distance between T361 and F389 during the same interval. **g** Distribution of the minimum distance of F365 sampled at three different voltages for K^+^ and Na^+^. **h** The relationship between the number of ion permeations per 20 ns interval and the corresponding average minimum distances between opposing E365 at the three different voltages. **i** A model illustrating the atomistic mechanism of voltage sensitivity in the pore domain.

Since the gate residues themselves do not possess elementary charges, we next questioned where the pore’s voltage sensitivity originates. To investigate this, we first calculated the backbone *φ* (phi) and ψ (psi) angle distribution sampled during the MD simulations at -100 mV and -150 mV for each residue in the pore, then derived the overlap of both *φ* and ψ angle distributions between the simulations performed at these two voltages (Fig. 7c). A high overlap, close to 1, indicates that the backbone conformation of the respective residue is highly similar between the two voltage conditions, while a low overlap indicates a voltage-dependent conformational change. The analysis indicated the largest conformational differences in two SF residues, I363 and G364 (Fig. 7c, d). Notably, G364 is located next to the acidic residue E365, which is located at the extracellular entrance of the SF and carries one elementary charge upon deprotonation. While the voltage-dependent backbone conformational change of E365 is rather minor (Fig. 7c), the side chain of E365 exhibits significant mobility during the MD simulations (Supplementary Fig. S2). We therefore computed the minimum distances of the two opposing E365 within 20 ns interval under three different transmembrane voltages and correlated them with the number of ion permeation during the same interval (Fig. 7g, h). As expected, larger E365 distances were observed at higher transmembrane voltages, and these were positively correlated with instantaneous ion conductance.

To further assess the impact of the protonation state of E365 on the pore’s voltage sensitivity, we conducted two sets of K^+^ permeation simulations at two different voltages, with (i) two of the four diagonally positioned E365 residues protonated, and (ii) all four E365 residues protonated (Fig. 4b). Overall, the simulations with two and four protonated E365 showed higher conductance compared to the deprotonated form, likely due to reduced interaction between uncharged E365s and the ions, which lowers the energy barrier for cation permeation. More importantly, these simulations revealed significantly reduced voltage sensitivity in the two protonated E365 configuration, while in the four E365 protonated form, voltage sensitivity was completely abolished.

Taken together, MD simulations of the CNGA1 pore revealed voltage-dependent conformational changes in the SF that correlate with the distance between opposing gate residues, thereby influencing ion conductance. We then explored how the SF and gate residues interact. A comparison of the open and closed state CNGA1 cryo-EM structures revealed a methyl-*π* interaction between T361 and F389 in the open state (Fig. 7e), which is absent in the closed state when F389 is oriented towards the pore. Since T361 is located directly adjacent to the intracellular side of the SF, it is reasonable to assume that this interaction mediates the communication between the SF and the gate. To further evaluate this hypothesis, we plotted the minimum distance between T361 and F389 within each 20 ns simulation interval against the number of ion permeation events during the same time interval. This analysis revealed a notable negative correlation (Fig. 7f): as the distance increased, indicating the loss of the interaction between T361 and F389, the number of ion permeation events decreased. These findings underscore the importance of the T361-F389 interaction in stabilizing the open state of the CNGA1 channel and mediating the communication between the SF and the gate (Fig.7i).

## Discussion

In this study, we conducted microsecond atomistic MD simulations to investigate the permeation of monovalent cations in the human CNGA1 channel. To the best of our knowledge, this is the first MD study to address ion permeation and conductance in native CNG channels. Leveraging the recent high-resolution cryo-EM structures of the open and closed states of the CNGA1 channel^32^, our study provides key atomistic mechanistic insights into the dynamics of ion permeation that are not accessible through the analysis of static cryo-EM structures.

By analyzing the ion binding profiles within the SF, we identified similar ion-binding sites but observed additional pronounced cation binding on the extracellular side of the SF, enabled by the negatively charged, mobile side chain of E365. This extracellular monovalent cation binding site was not identified in the cryo-EM data. We hypothesize that this is due to the high dynamics of this binding site or the protonation of E365 under the physiological conditions. Since the overall dimensions of the SF in the CNGA1 channel are much wider than those of typical K^+^ channels, cation binding in the SF region is generally more diffusive than in K^+^ channels^56, 57^. Consequently, we conclude that the SF of CNGA1 does not present a high energy barrier for monovalent cation conduction. Furthermore, the wide SF in the CNGA1 channel allows it to accommodate both Na^+^ and K^+^ in their partially hydrated forms, without requiring the additional conformational changes observed in some other non-selective cation channels, such as the NaK channel^52^ and the lysosomal two pore channel 2 (TPC2)^58^.

In contrast to many other computational investigations of ion channels that employed substantially larger transmembrane voltages (300-600 mV) to accelerate the ion permeation process, we used here a range of transmembrane voltages close to those in experimental electrophysiology studies. For comparison, we also performed ion permeation simulations of CNGA1 with the AMBER99SB and AMBER19SB force fields. Overall, we observed lower conductance in the AMBER simulations, particularly an unrealistically low Na^+^ conductance (Supplementary Table S1), which was also seen in our previous simulations with the AMPA GluA2 receptor ion channel^55^. Notably, the simulated K^+^ conductance of the pore domain and the VSD using the CHARMM36m force field closely matches single-channel recording data at +100 mV and at saturating cGMP concentration^10^. The simulated Na^+^ conductance is notably higher than the K^+^ conductance, which also qualitatively aligns with experimental observations. From the MD simulations, we proposed that the higher Na^+^ density in the pore, rather than the previously assumed shorter dwell time of Na^+^, is the primary mechanism underlying the larger Na^+^ single-channel conductance than K^+^.

A further main finding of this computational investigation of the CNGA1 is a plausible explanation for the excess noise of the single-channel conductance observed experimentally in the open state. In addition to the previously assumed fluctuation of the pore, we revealed that the changes in hydration level in the pore, caused by the fluctuation of two large hydrophobic gate residues, F389 and V393, highly influence the ion conduction rate. Furthermore, from the simulations, we propose that the dehydration of the gate could rapidly lead the channel into a non-conductive state without the need for a full transition into the closed state. This may provide additional insights into the rapid flickery gating behavior characteristic of CNG channels^59, 60^. It is worth noting that hydrophobic gating has been previously proposed by MD simulations in various ion channels^61^.

Lastly, by simulating the CNGA1 channel pore domain alone, we observed an unexpected voltage sensitivity in its conductance. This voltage sensitivity diminished when the gate was restrained or when the acidic E365 residues were protonated throughout the simulations. We are aware that protonation and deprotonation are transient processes, and in narrow pore regions, two nearby titratable residues could even share a proton, as observed in the M2 channel^62^. However, accurately modeling proton dynamics is challenging in classical MD simulations, where future studies using quantum-mechanics/molecular mechanics (QM/MM) approaches are needed. Our current simulations, which vary the number of protonated E365, are limited to providing insights into how the electrostatic effects resulting from deprotonation influence ion conductance and voltage sensitivity. Moreover, a detailed analysis of the MD data suggested a coupling mechanism between the SF and gate residue F389, mediated by the interaction of F389 and T361, a residue adjacent to the SF (Fig. 7g). Interestingly, previous atomistic MD simulations of K^+^ channels have highlighted the importance of similar interactions, such as T59-I84 in MthK^63^ and T280-F316 in TREK channels^64^, in facilitating communication talk between the selectivity gate and the pore-lining helices. Therefore, experimentally mutating threonine to serine at residue 361 would be a useful approach to validate our simulation results, as we expect that the absence of the methyl group in the serine side chain would reduce the cross-talk between the SF and the gate.

In addition, we performed a protein structure network analysis using PSNTool^65, 66^ based on the MD trajectory to identify the shortest prevalent communication pathways in the CNGA1 channel. In the simulations of the pore domain only, the major identified pathways highlighted the communication between the gate and the SF (Supplementary Fig. S6). In sharp contrast, for the simulations that included both the pore domain and the VSD, where the voltage-dependent conductance behavior was absent, the interactions between the VSD and the pore domain became the main interaction pathways (Supplementary Fig. S6). This result indicates that the voltage sensitivity in the pore, enabled by the flexibility of the pore, diminishes through the interaction of the pore domain with the VSD. In general, this finding aligns with a previous mutation study of the CNGA1 channel, which indicated that proper attachment of the SF to the surrounding channel pore is essential to minimize voltage gating in CNG channels^16^. Despite being named the voltage-sensing domain, the VSD in the CNG channel is not responsive to voltage changes due to the unique location of a series of positively charged residues in the VSD that are outside the transmembrane voltage field^29, 32^. Ironically, the VSD rather acts as the voltage-stabilizing domain in CNGA1, abolishing the voltage sensitivity of the pore enabled by its tight interaction with the pore.

In conclusion, large-scale atomistic MD simulations of ion permeation in the CNGA1 channel confirmed that the open state of CNGA1, as resolved by cryo-EM, is the main conductive state. This finding is supported by the simulated K^+^ conductance matching the experimental single-channel conductance at saturating cGMP concentrations. We demonstrated that monovalent cations traverse the pore by primarily binding in the SF region in a partially dehydrated form for both K^+^ and Na^+^, providing atomistic insights into cation non-selectivity. Additionally, we identified a hydrophobic funnel in the central gate cavity that significantly influences ion conductance, potentially explaining the flickery gating behavior and excess single-channel conductance noise observed experimentally. Finally, we identified an unexpected voltage-dependent conductance when only the pore was simulated, driven by long-range coupling between the deprotonated E365 in the SF and the gate. This voltage sensitivity was abolished when pore flexibility was reduced or when the E365 residues were protonated. These findings offer atomistic insights into the voltage sensitivity of the pore in CNG channel, a phenomenon observed in decades of electrophysiological studies but previously elusive at the atomistic level.

## Methods

### Atomistic molecular dynamics simulation

In this study, MD simulations started from the high-resolution cryo-EM structure of the homomeric CNGA1 channel in the open state (PDB ID: 7LFW) and the closed state (PDB ID: 7LFT)^32^. The CNGA1 channel was simulated using two different structural models: the full transmembrane domain, encompassing both the pore domain and the voltage-sensing domain (VSD) (Fig. 4a); an isolated pore domain model (Fig. 4b, c), a strategy frequently employed in MD simulations of ion channels to mitigate computational complexity^53, 54, 55^. Since we were not simulating the full-length construct and to prevent any artifacts originating from charges at the N- and C-termini, the structures were neutralized by adding acetyl and N-methyl caps using PyMOL^67^ Builder option.

The simulations were mainly performed using the CHARMM36m force field^40^, while several simulations were conducted with the AMBER99SB^41^ and AMBER19SB^42^ force fields for validation purposes (Supplementary Table S1). The CNGA1 channel structures were embedded within a palmitoyloleoyl phosphatidylcholine (POPC) lipid bilayer. In the AMBER99SB simulation setup, we inserted the CNGA1 channel into the POPC lipid bilayer with the GROMACS internal embedding function. Improved ion parameters^68^ and lipid parameters^69^ were employed in these simulations. The insertion of CNGA1 channel into the lipid bilayer was carried out in CHARMM-GUI^70^ for AMBER19SB and CHARMM36m simulations. TIP3P (Transferable Intermolecular Potential with Three Points) water model^71^ was used in all simulations. The concentrations of KCl and NaCl were set to 150 mM.

After the CNGA1 channel was embedded into a POPC lipid bilayer with ions and water in a box, the simulation system was energy minimized and equilibrated. The energy minimization was done with GROMACS ‘steepest descent’ algorithm to reduce the system maximum force to below 1000 kJ/mol/nm in 50,000 steps. After the energy minimization, a 10 ns equilibration without any restraints was performed. For the simulations prepared by the CHARMM-GUI, the system was energy minimized and equilibrated in six steps using default scripts provided by the CHARMM-GUI.

For all simulation of the CNGA1 pore domain using CHARMM36m force field and all simulations of CNGA1 pore domain together with VSD, the transmembrane electric potential was directly generated by an external electric field applied along the z-axis (pore axis) (Supplementary Table S1). The voltage was calculated with:

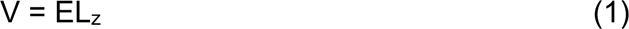

where E denotes the applied electric field and L_z_ is the length of the simulation box along the z-axis^36, 72, 73^. The first 100 ns of these simulations (50 ns in closed structure simulations) were considered as equilibration phases and discarded for the further analysis (Supplementary Fig. S7).

For comparison purposes, for some simulations of the CNGA1 pore domain using AMBER force field, we used computational electrophysiology (CompEL) setup (Supplementary Table S1)^74^. To establish a transmembrane potential, a copy of the equilibrated system was stacked along the pore axis, generating two compartments (an inner and an outer). After building the two-bilayers system, we conducted another round of equilibration for 20 ns without cation imbalance before the actual MD production run. For the production run, a charge imbalance of 2 *e* between the two compartments separated by two lipid bilayers resulted in transmembrane voltages that are listed in Supplementary Table S1. During the MD simulations, ions passing through the pore were regularly monitored, and the charge imbalance was maintained by exchanging the same species of ion in one compartment with a water molecule from the other compartment^74^. The resulting transmembrane voltage can be calculated by double integration of the charge distribution using the Poisson equation as implemented in the GROMACS tool *gmx potential^75^*.

All atomistic molecular dynamics simulations were run on GROMACS software package (version 2021.1 and 2023.3)^76^. Short-range electrostatic interactions were calculated with a cutoff of 1.0 nm, whereas the long-range electrostatic interactions were treated using the particle mesh Ewald method ^77^. The cutoff for van der Waals interaction was set to 1.0 nm. The simulations were performed at 300 K with an enhanced Berendsen thermostat (GROMACS V-rescale thermostat^78^. The Parrinello-Rahman barostat^79^ was employed to keep the pressure within the system remaining at 1 bar. All bonds were constrained with the Linear Constraint Solver (LINCS) algorithm^80^. We used virtual sites approach to reduce the computational cost when adopting Amber99SB force field, which allowed us to increase the integration time step to 4 fs. Otherwise, all simulations were performed with an integration time step of 2 fs.

The protonation states of all titratable residues were assigned according to the standard protonation states at pH 7. Systems with varied protonation states at E365 were prepared by using CHARMM-GUI webserver tool^70^.

To investigate the influence of gate distance on the conductance, we applied harmonic distance restraints between the C*α* atoms of the gate residue F389. We used the distances determined from the Cryo-EM structure and applied the distance restraints to both the opposite and adjacent subunits. Two distance restraints were applied on the opposite C*α* of F389 with 1.93 nm and four distance restraints were applied on the adjacent C*α* of F389 with 1.38 nm. The distance restraint force constant for all restraints was set to 1000 kJ/mol/nm^2^.

All trajectories were analyzed with GROMACS toolkits and Python3 using MDAnalysis^81^ together with the package Numpy^82^, Matplotlib^83^, Pandas^84^ and SciPy^85^. Residue-residue pair distances were calculated using GROMACS tool *gmx dist* and dihedral angles were calculated with *gmx rama*. Presented data of dihedral angle distribution and residue-residue pair distance distribution were calculated using kernel density estimation^86^ from five parallel simulation runs. Ion permeation events were calculated when a cation permeated through the entire pore from the SF to the gate. To determine the ion hydration states during permeation, we calculated the number of ion-coordinating oxygens from both water molecules and protein residues within each ion’s first solvation shell. Hydration shells are dynamic; here, we defined the waters of hydration using the radii corresponding to the minimum in the radius of gyration profiles: 3.1 Å for Na^+^ and 3.4 Å for K^+87^. Instantaneous conductance was calculated by counting ion permeation events over 20 ns from five independent simulations. An ion permeation event was counted if the ion passed through the gate. Molecular visualizations were made with PyMol and Visual Molecular Dynamics (VMD)^88^.

Protein structure network analysis was conducted using PSNTool^65, 66^ based on the MD trajectories from five parallel production runs. A graph was defined by a set of vertices (nodes) and connections (links) between them. In a PSN, each amino acid residue was represented as a node and these nodes were connected by links based on the strength of non-covalent interactions between residues. A global metapath composed of the most recurrent links in the shortest path pool was computed to infer a coarse picture of the structural communication.

All simulation details were summarized in Supplementary Table S1 and S2.

## Data availability

All data that support the findings of this study are included in this manuscript and in the Supplementary Information. All source data, Supplementary Movies and the simulation run input files, comprising the starting configuration and all necessary parameters for performing the MD simulations are deposited in Zenodo under accession code https://doi.org/10.5281/zenodo.14508687

## Code availability

Molecular dynamics simulation data were generated using GROMACS 2021.1/2023.3. All trajectories were analyzed with GROMACS tools and Python using MDAnalysis together with Numpy, Matplotlib, Pandas and SciPy.

## Acknowledgements

This work was funded by the Leibniz-Forschungsinstitut für Molekulare Pharmakologie (FMP) and Deutsche Forschungsgemeinschaft (DFG) CRC1078 ‘Protonation Dynamics in Protein Function’ and RU2518 DynIon (to H.S.). The MD simulations were performed with resources provided by the North-German Supercomputing Alliance (HLRN), the high-performance computer “Lise” at the NHR Center (NHR@ZIB) and Erlangen National High Performance Computing Center (NHR@FAU). We thank Prof. Klaus Benndorf, Dr. Jana Kusch, and Prof. Andrew Plested for helpful discussions. We would like to acknowledge the technical support of Dr. Songhwan Hwang and Dr. Tillmann Utesch.

## Author contributions

H.S. and H.L. designed and directed the project. H.L. performed the MD simulations.

H.L. analyzed the data with the contributions from J.B. The manuscript was written by H.L. and H.S.

## Supplementary Information

### Supplementary Figures

**Supplementary Fig. S1.**
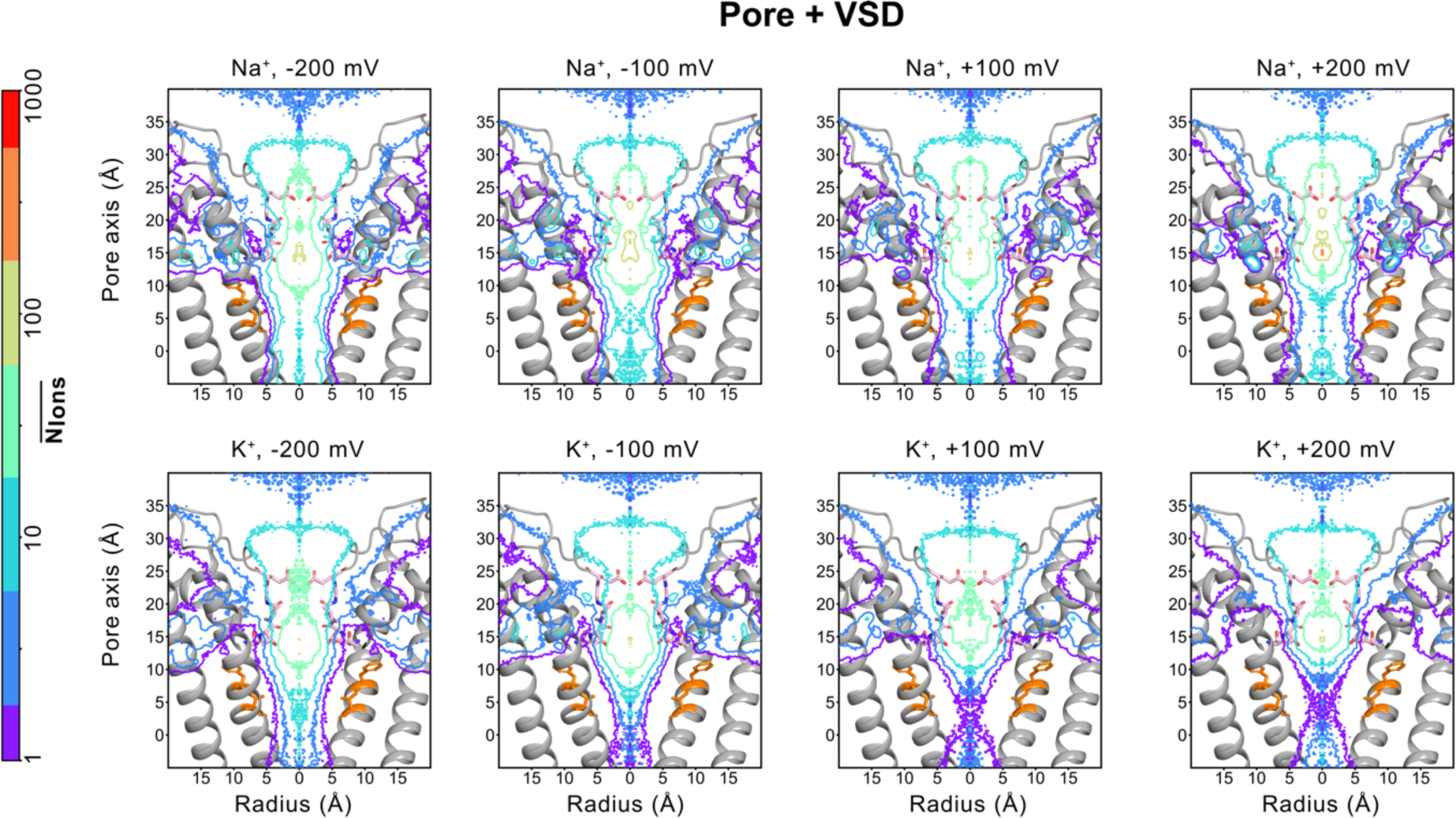
Ion binding profiles of K^+^ and Na^+^ within the ion conduction pathway derived from MD simulations of the pore domain together with the voltage-sensor domain (VSD). Comparison of 2-dimensional ion occupancy maps for the CNGA1 channel under varying voltage conditions with Na^+^ and K^+^. The simulations were performed with the pore domain together with the voltage-sensor domain (VSD).

**Supplementary Fig. S2.**
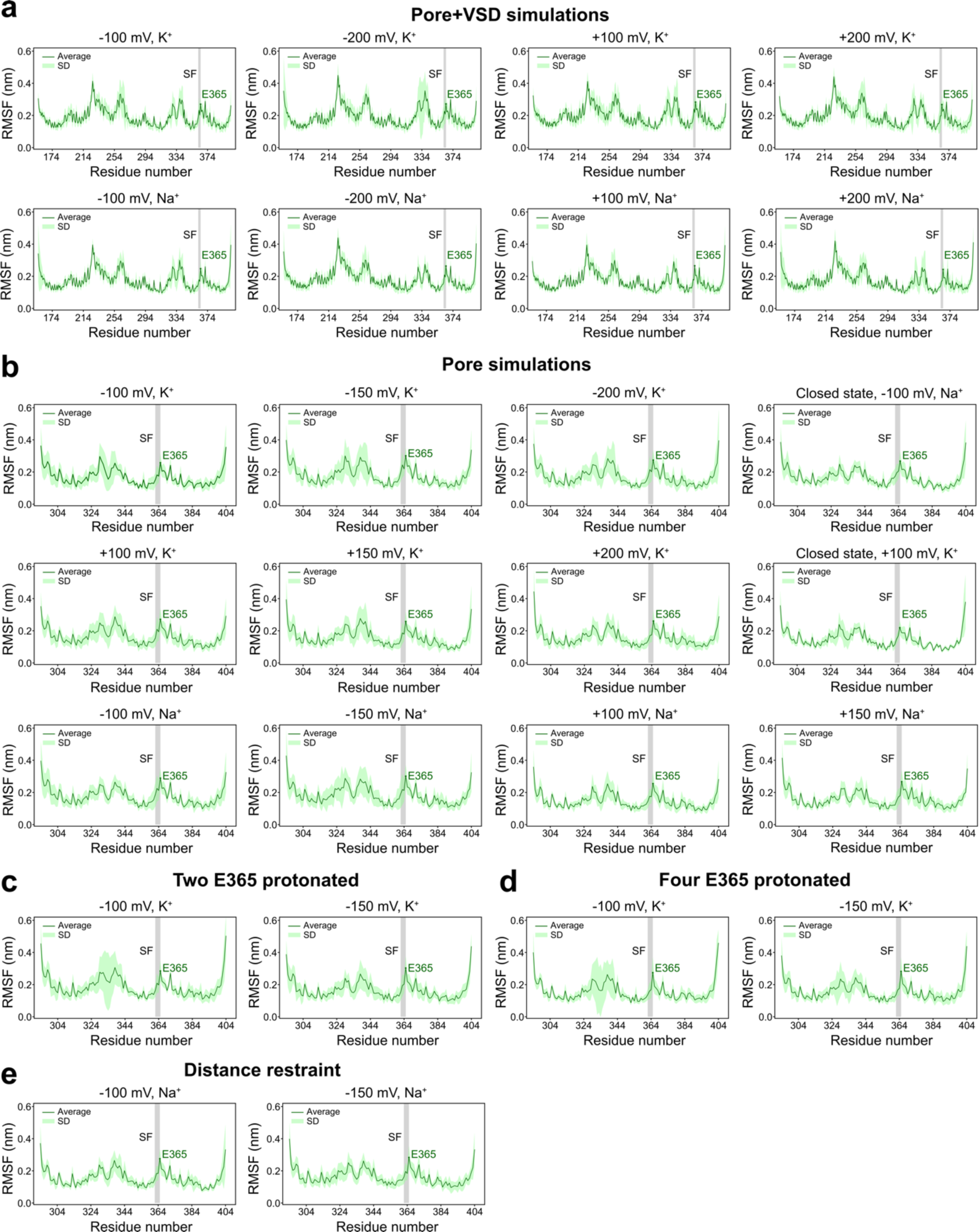
Root-mean-square-fluctuation (RMSF) of the MD simulations. For each MD simulation setup, the green line represents the average RMSF value from five parallel runs, while the light green shading indicates the standard deviation. The grey-highlighted region marks the selectivity filter (SF), with E365 showing increased dynamics. **a** Simulations of the pore domain and the voltage-sensor domain (VSD) of the CNGA1 channel with K^+^ and Na^+^ under various transmembrane voltages. **b** Simulations of only the pore domain of the CNGA1 channel with K^+^ and Na^+^ under various transmembrane voltages. **c** Simulations of the pore domain of the CNGA1 channel, where two opposing E365 residues are protonated, with K^+^ under -100 mV and -150 mV voltages. **d** Simulations of the pore domain of the CNGA1 channel, where all four E365 residues are protonated, with K^+^ under -100 mV and -150 mV voltages. **e** Simulations of the pore domain of the CNGA1 channel with distance restraints on gate residue F389 with Na^+^ under -100 mV and - 150 mV voltages.

**Supplementary Fig. S3.**
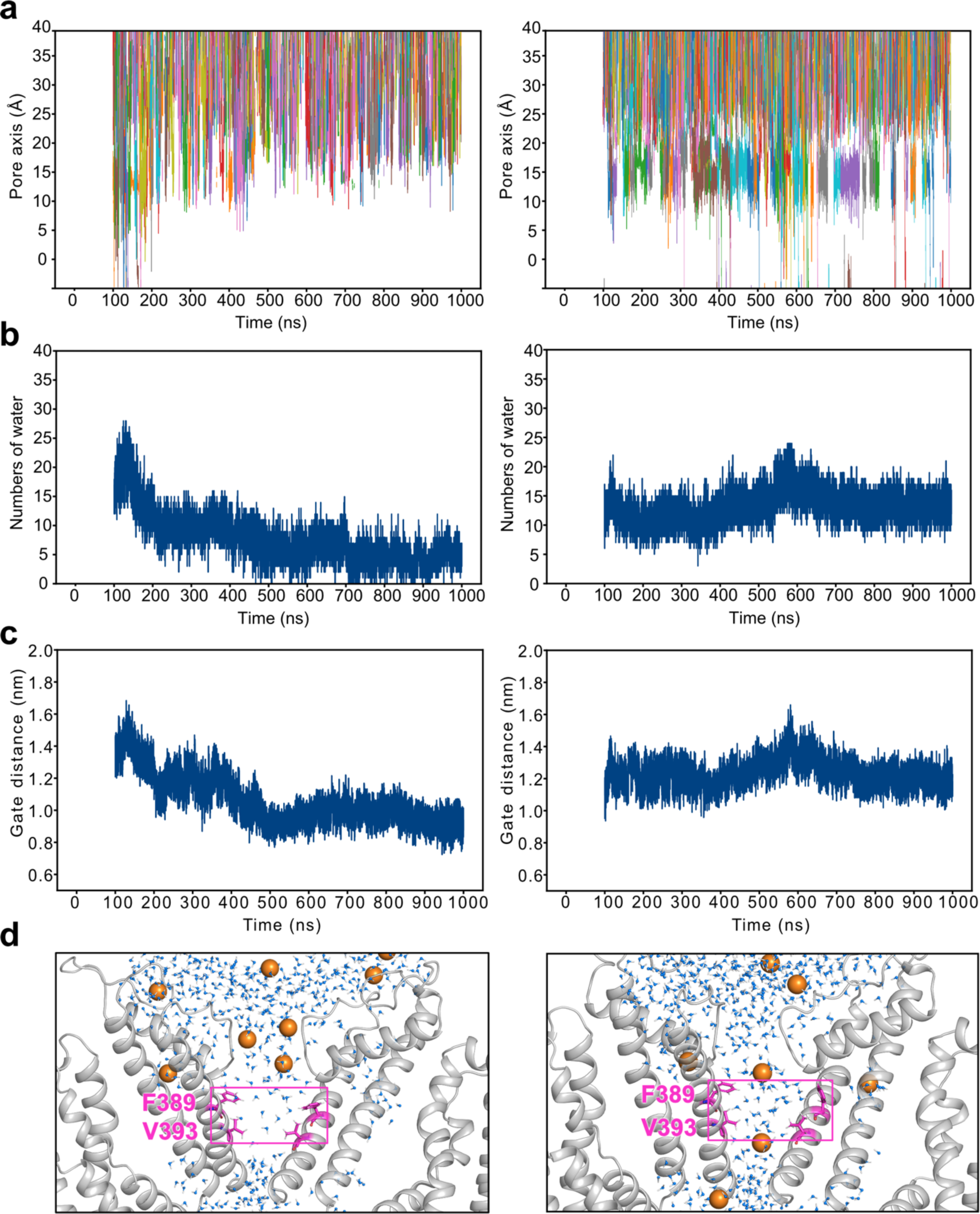
Examples of ion permeations stopping due to partial gate closure and dehydration. **a** Representative traces of K^+^ ions passing through the pore of the CNGA1 channel. The left panel shows a trajectory where ion flow stops after 100 ns, while the right panel shows continuous ion permeations throughout the simulations. **b** The number of water molecules in the gate region over time during the simulations shown in (**a**). **c** The distance between two opposing gate residues F389 over time during the simulations in (**a**). **d** Two representative snapshots showing the hydration level differences in the gate region: the left panel shows partial dehydration, and the right panel shows full hydration.

**Supplementary Fig. S4.**
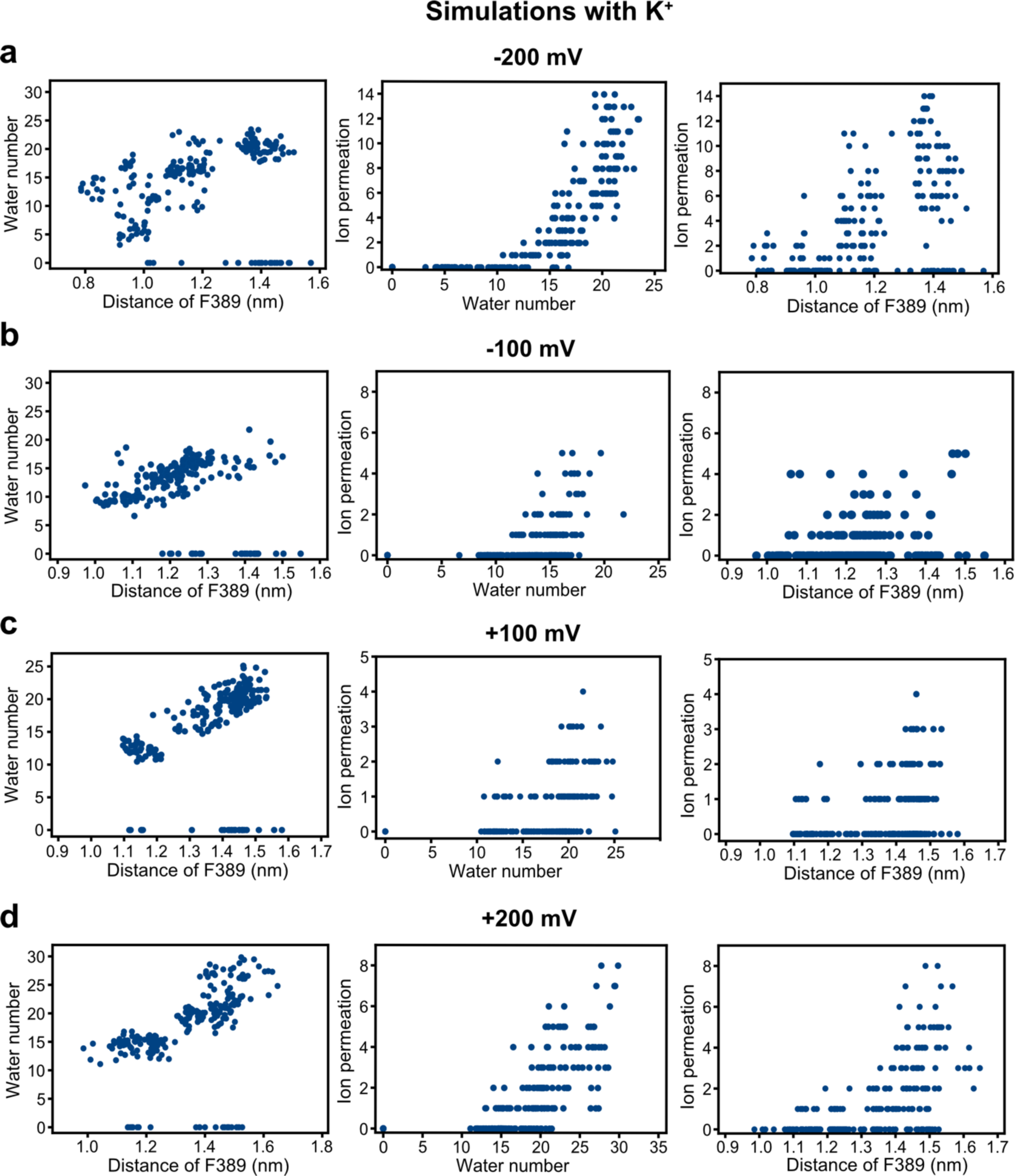
The relationship between the average number of water molecules in the gate region per 20 ns interval, the average distance between the two opposing gate residues (F389), and the number of ion permeation events during the same time interval. K^+^ permeation simulations were performed at -200 mV, -100 mV, +100 mV, and +200 mV, respectively. The simulations were performed with the pore domain together with the voltage-sensor domain (VSD).

**Supplementary Fig. S5.**
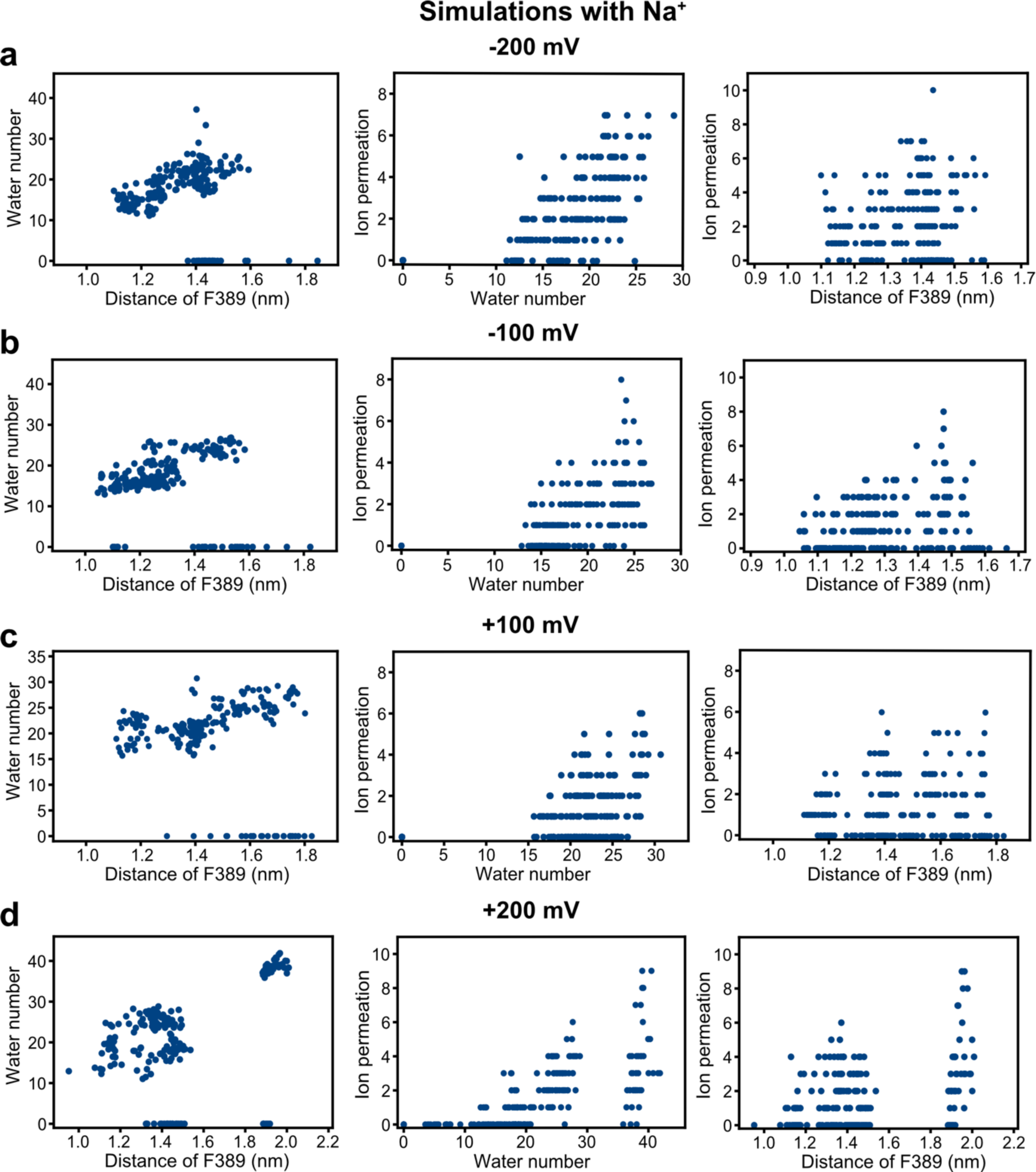
The relationship between the average number of water molecules in the gate region per 20 ns interval, the average distance between the two opposing gate residues (F389), and the number of ion permeation events during the same time interval. Na^+^ permeation simulations were performed at -200 mV, -100 mV, +100 mV, and +200 mV, respectively. The simulations were performed with the pore domain together with the voltage-sensor domain (VSD).

**Supplementary Fig. S6.**
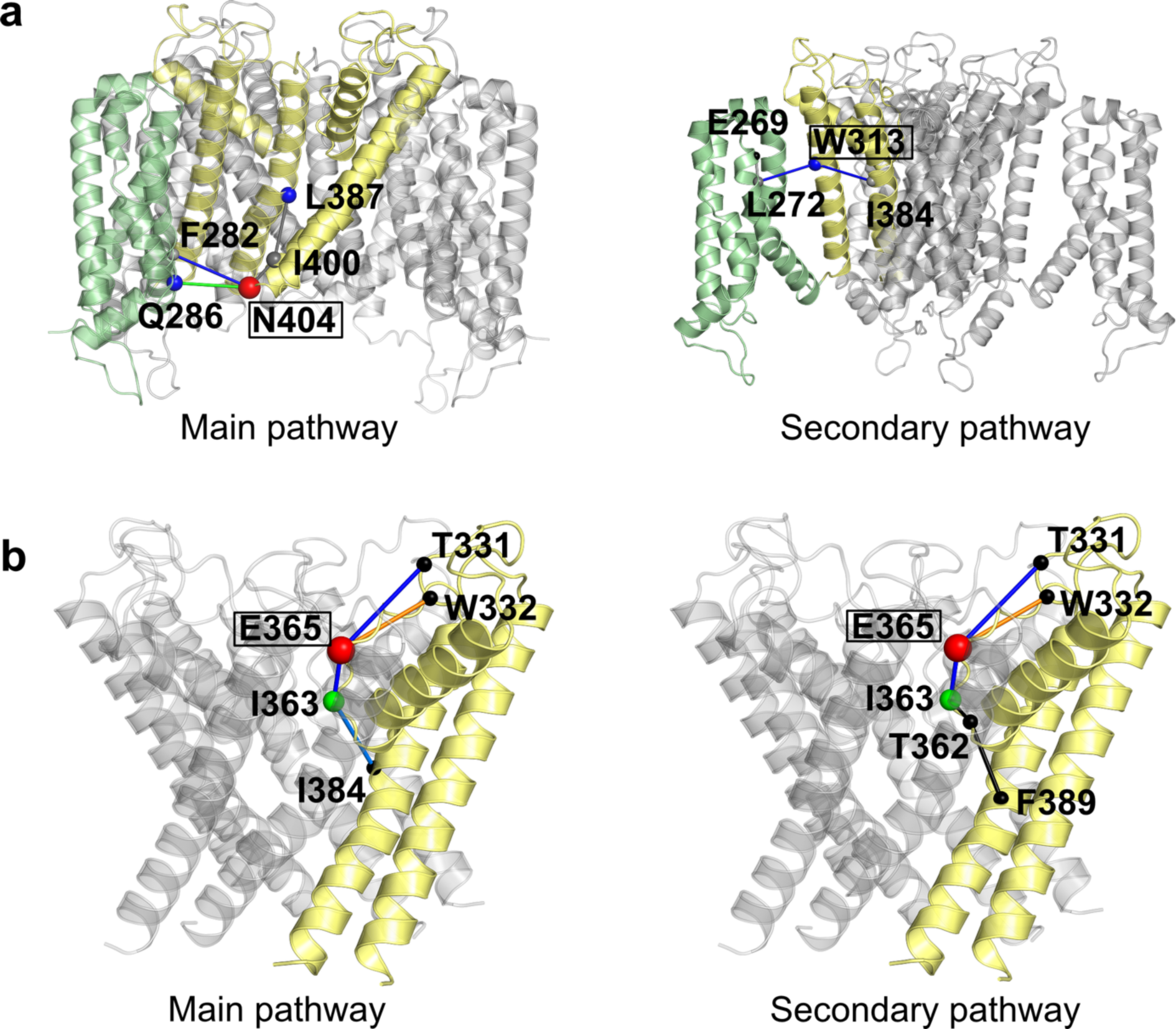
Protein structure network analysis of the CNGA1 channel using PSNTool^1, 2^. **a** Primary and secondary protein structure pathways in simulations of pore domain and VSD. N404 and W313 were revealed to be the most important connection node residues in their respective pathways. **b** Primary and secondary protein structure pathways of simulations involving only the pore domain. E365 was revealed to be the most important connection node residue in both pathways. Connection node residues are represented as spheres, with interactions depicted as links. The size of the sphere reflects the interaction frequency of the residues, with larger spheres indicating more connections to other residues. The color of the sphere and link represents the interaction strength (red > orange > green > blue > grey > black).

**Supplementary Fig. S7.**
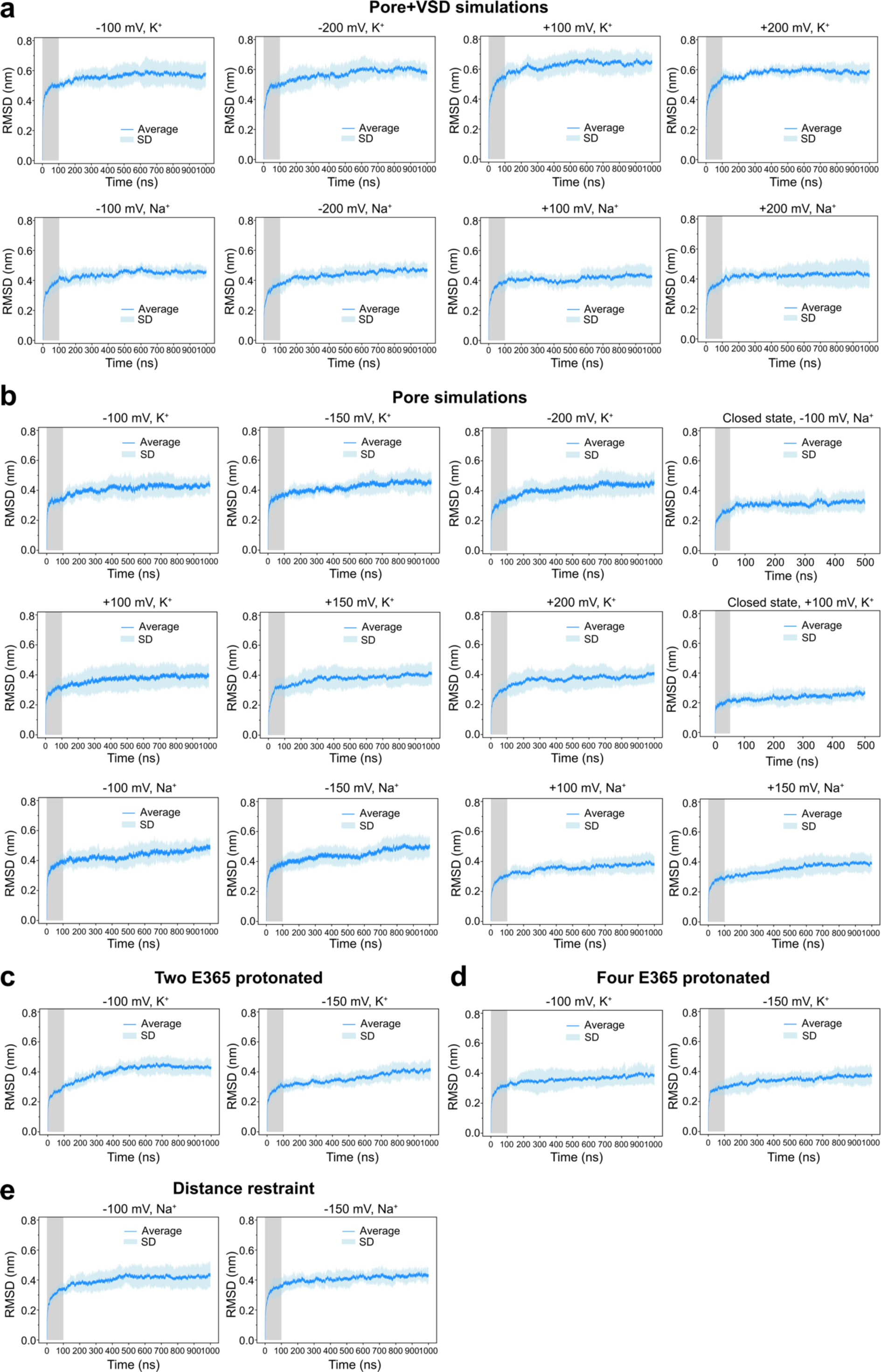
Root-mean-square-deviation (RMSD) of the MD simulations. For each MD simulation setup, the blue line represents the average RMSD value from five parallel runs, with the blue shading indicating the standard deviation. The grey-shaded area illustrates the equilibration period for each simulation: 100 ns for 1 μs and 50 ns for the 500 ns simulations. The equilibration periods were excluded from the subsequent analyses. **a** Simulations involving the pore domain and voltage-sensor domain (VSD) of the CNGA1 channel with K^+^ and Na^+^ under various transmembrane voltages. **b** Simulations involving only the pore domain of the CNGA1 channel with K^+^ and Na^+^ under various transmembrane voltages. **c** Simulations of the pore domain of the CNGA1 channel, where two opposing E365 residues are protonated, with K^+^ under -100 mV and -150 mV voltages. **d** Simulations of the pore domain of the CNGA1 channel, where all four E365 residues are protonated, with K^+^ under -100 mV and -150 mV voltages. **(e)** Simulations of the pore domain of the CNGA1 channel with distance restraints on gate residues F389 with Na^+^ under -100 mV and -150 mV voltages.

### Supplementary Tables

**Supplementary Table S1.**
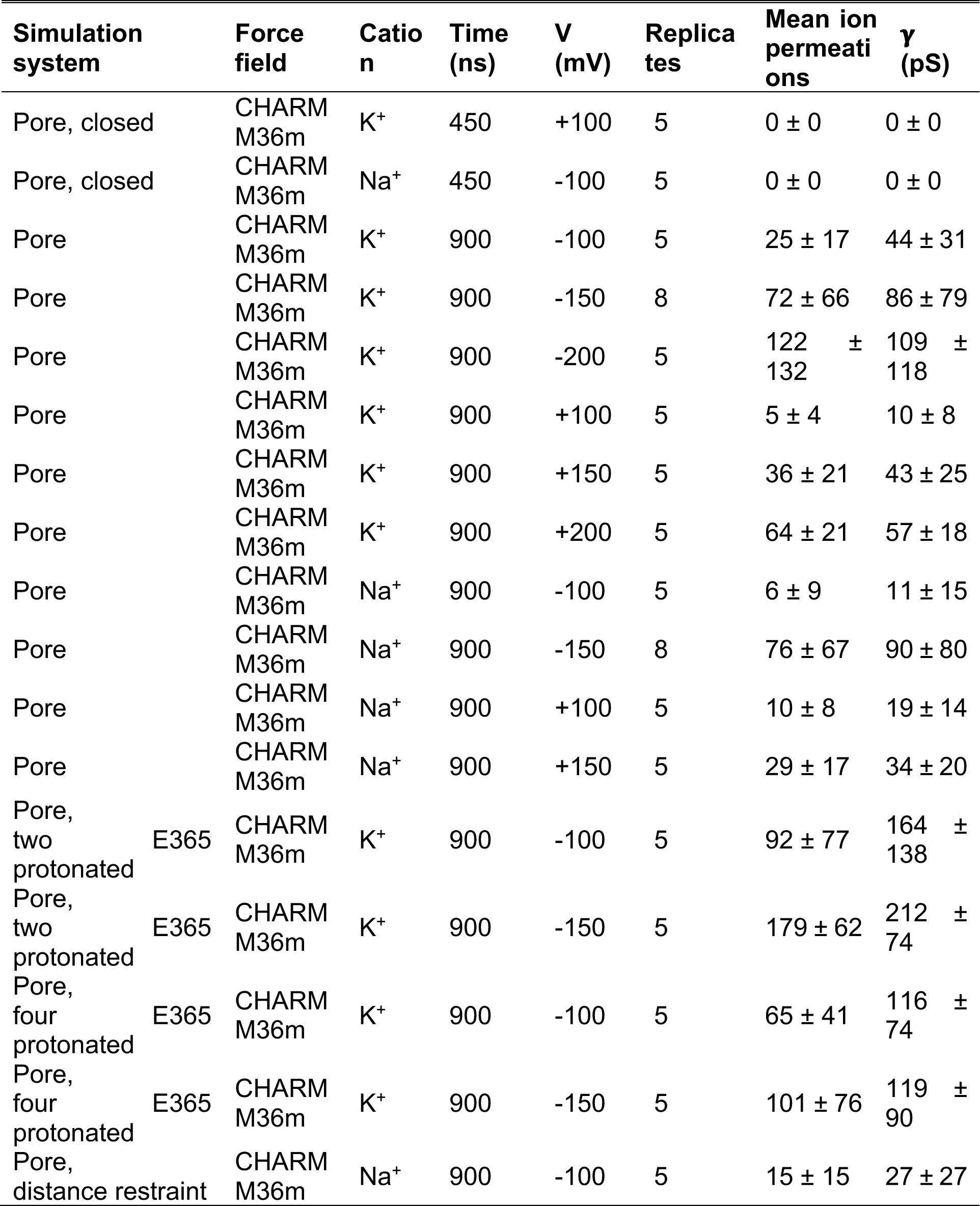
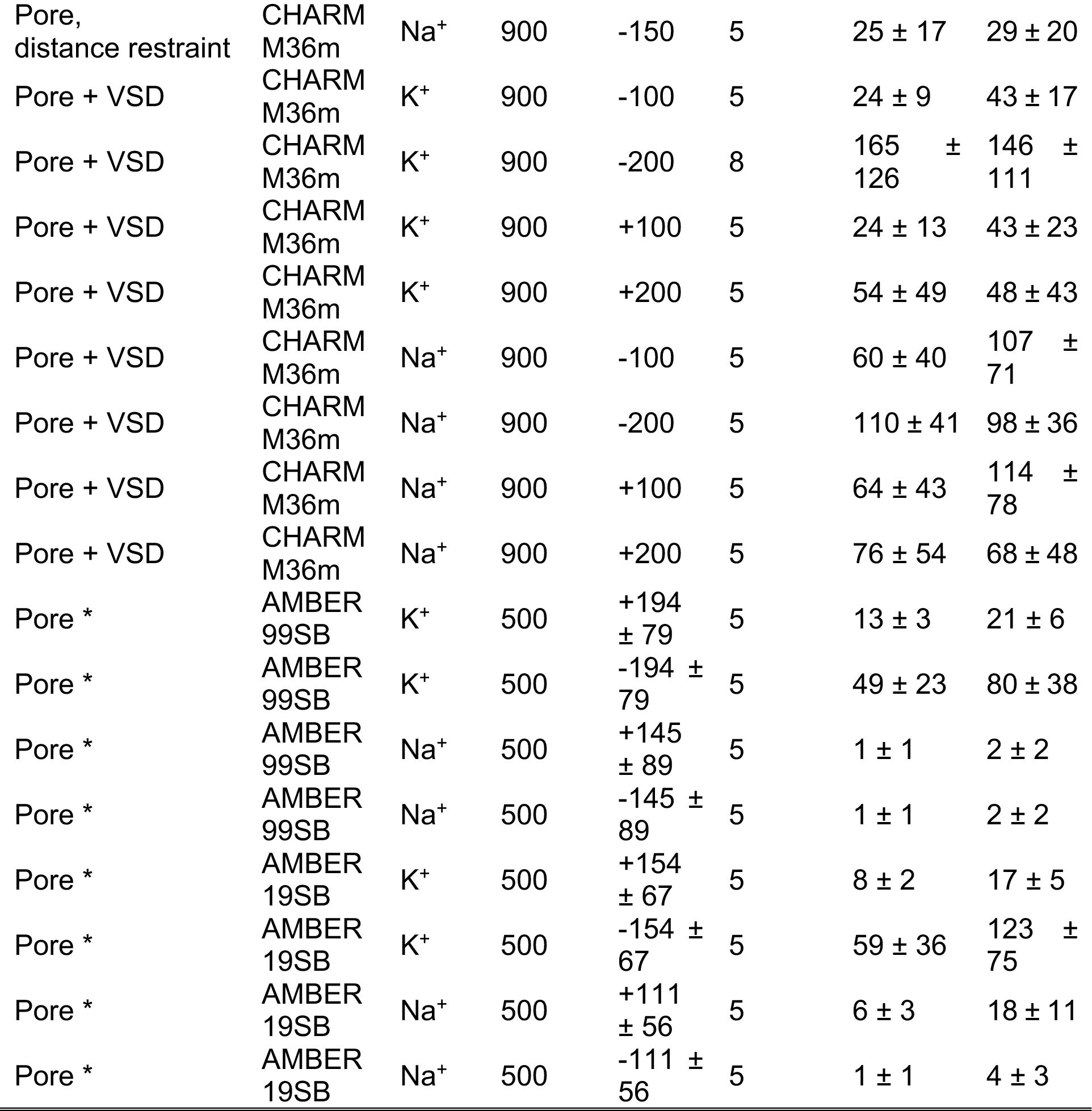
List of simulation systems. Simulations involving the CNGA1 channel pore domain or the pore domain together with the VSD for Na^+^ and K^+^ under different voltages. All of the simulations were performed with a salt concentration of 150 mM and a temperature at 300 K. * Simulations using computational electrophysiology method^3^; others using an external electric field.

**Supplementary Table S2.**
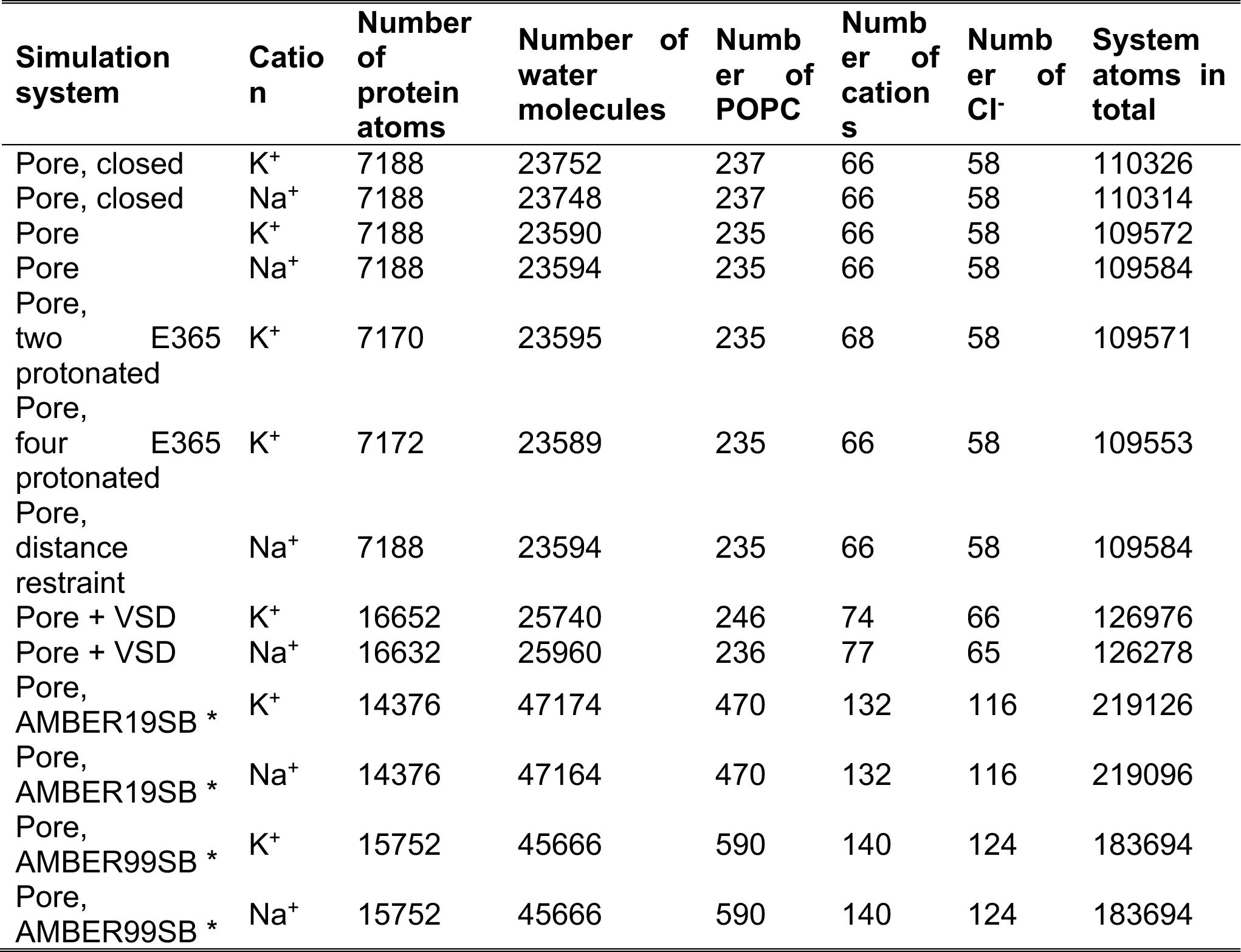
Simulation system details. The simulations without special indication were performed with the CHARMM36m force field. * Simulations using computational electrophysiology method^3^; others using an external electric field.

### Supplementary Movies

**Supplementary Movie S1 | K^+^ permeation in CNGA1 channel during a 50 ns trajectory.**

This simulation was performed with only the pore domain under an external electric field of -150 mV. For clarity, only two subunits of CNGA1 were shown in cartoon (silver), the SF is shown in magenta sticks, and the gate in orange sticks. Permeating K^+^ ions are shown as spheres of various colors, while the rest of the K^+^ ions are shown in black. Water molecules are drawn as blue O bonded to white H atoms. The Z-positions of permeating K^+^ ions along the pore axis are monitored during ion permeation.

**Supplementary Movie S2 | Na^+^ permeation in CNGA1 channel during a 40 ns trajectory.** This simulation was performed with only the pore domain under an external electric field of -150 mV. For clarity, only two subunits of CNGA1 were shown in cartoon (silver). Permeating Na^+^ ions are shown as spheres of various colors, while the rest of the Na^+^ ions are shown in black. Water molecules are drawn as blue O bonded to white H atoms. The Z-positions of permeating Na^+^ ions along the pore axis are monitored during ion permeation.

